# Neural computations in prosopagnosia

**DOI:** 10.1101/2022.12.13.519604

**Authors:** Simon Faghel-Soubeyrand, Anne-Raphaelle Richoz, Delphine Waeber, Jessica Woodhams, Frédéric Gosselin, Roberto Caldara, Ian Charest

## Abstract

We aimed to identify neural computations underlying the loss of face identification ability by modelling the brain activity of brain-lesioned patient PS, a well-documented case of acquired pure prosopagnosia. We collected a large dataset of high-density electrophysiological (EEG) recordings from PS and neurotypicals while they completed a one-back task on a stream of face, object, animal and scene images. We found reduced neural decoding of face identity around the N170 window in PS, and conjointly revealed normal *non-face* identification in this patient. We used Representational Similarity Analysis (RSA) to correlate human EEG representations with those of deep neural network (DNN) models of vision and caption-level semantics, offering a window into the neural computations at play in patient PS’s deficits. Brain representational dissimilarity matrices (RDMs) were computed for each participant at 4 ms steps using cross-validated classifiers. PS’s brain RDMs showed significant reliability across sessions, indicating meaningful measurements of brain representations with RSA even in the presence of significant lesions. Crucially, computational analyses were able to reveal PS’s representational deficits in high-level visual and semantic brain computations. Such multi-modal data-driven characterisations of prosopagnosia highlight the complex nature of processes contributing to face recognition in the human brain.

**Highlights:**

- We assess the neural computations in the prosopagnosic patient PS using EEG, RSA, and deep neural networks
- Neural dynamics of brain-lesioned PS are reliably captured using RSA
- Neural decoding shows normal evidence for non-face individuation in PS
- Neural decoding shows abnormal neural evidence for face individuation in PS
- PS shows impaired high-level visual and semantic neural computations

## Introduction

The human brain is equipped with sophisticated machinery optimised to quickly and effectively recognise faces in a series of computations unfolding within tens of milliseconds. A dramatic contrast to this typically efficient process has been revealed in brain-lesioned patients with an inability to recognise faces, individuals called acquired prosopagnosics (Bodamer, 1947). Findings from these patients have refined the functional role and the distributed nature of the face-sensitive brain regions in the ventral stream, such as the fusiform gyrus (FFA; Bobes et al., 2003; Kanwisher et al., 1997) and the lateral portion of the inferior occipital gyrus (Occipital Face Area, OFA; Dricot et al., 2008; Gauthier et al., 2000; Rossion et al., 2003; Sorger et al., 2007). This literature has generally contributed to the idea that specialised and category-selective neural modules are necessary for functional aspects of face processing (Cohen et al., 2019). Brain imaging findings from individuals born with deficits in face recognition (developmental prosopagnosics; (Avidan et al., 2005; Jiahui et al., 2018; Kaltwasser et al., 2014; McConachie, 1976; Rosenthal et al., 2017) have revealed finer-grained functional neural differences in the processes (Jiahui et al. 2018; Rosenthal et al. 2017; Avidan et al. 2014; Zhao et al. 2018) associated with deficits in face recognition. Overall, the cumulation of these neuropsychological, neuroanatomical and functional components of prosopagnosia (Busigny et al. 2010; Dricot et al. 2008; Rossion 2018; Rossion et al. 2003; Duchaine and Nakayama 2006) has significantly contributed to neural models of face perception in the last two decades (Duchaine & Yovel, 2015; Haxby et al., 2000; White & Mike Burton, 2022). Yet, very little is known on the nature of face representations of those patients (e.g., Caldara et al., 2005; Fiset et al., 2017), and next to nothing is known on the nature of brain dynamics and neural computations affected in prosopagnosia. Here, we report an investigation of the neural computations involved in the processing of faces and objects of patient PS, a well-documented case of pure acquired prosopagnosia (Rossion et al., 2003; Sorger et al., 2007), using Representational Similarity Analysis (RSA) applied to brain imaging and computational models.

Patient PS is a right-handed woman having sustained a closed head injury in 1992, leading to extensive bilateral occipitotemporal lesions encompassing the right OFA, left FFA, and a small region of the right middle temporal gyrus (Dricot et al., 2008; Sorger et al., 2007). She is perhaps the most studied case of acquired prosopagnosia, with more than 32 scientific publications in the last 20 years (see Rossion, 2022a, 2022b for recent reviews on this patient). The particular attention given to this case can be attributed to the relatively focal aspect of her lesions in the face network and the resulting highly specific impairment for face-identification (Busigny et al., 2010). The neuro-anatomical/functional basis of PS has already been exhaustively reviewed elsewhere (Rossion, 2022b). Overall, while her condition has been shown to affect a wide array of perceptual mechanisms (e.g. holistic processes, Ramon et al., 2016; the visual content of face representations in Caldara et al., 2005; Fiset et al., 2017; see also Rossion, 2022a), a direct characterisation of the neural computations behind her deficits has, to the best of our knowledge, never been attempted. A traditional proxy to the nature and level of brain computations affected in this patient, and in prosopagnosia in general, has been to consider the temporal dynamics and face-selectivity of the underlying neural activity. Event-related potential differences occurring late, for example, are generally interpreted as representing higher level processes than those occurring earlier (Alonso Prieto et al., 2011; Bentin & Deouell, 2000; Eimer et al., 2012; Gosling & Eimer, 2011; Herzmann et al., 2004; Liu-Shuang et al., 2016a; Simon et al., 2011; Tanaka et al., 2006; Wiese et al., 2019a). Some associations have been revealed between prosopagnosia and typical neural correlates of face-processing, like the face-sensitive N170 (Bentin et al., 1996) and face-selective fMRI activation (Towler and Eimer 2012; Bobes et al. 2003; Alonso Prieto et al. 2011; Gao et al. 2019). Interestingly, however, despite important lesions and behavioural deficits in face-identification, PS still shows typical face-selectivity in spared-regions of the right-hemisphere (i.e. she displays a right FFA; e.g. see Gao et al., 2019; Rossion et al., 2003), as well as a typical N170 component in the right, but not left-hemisphere (Alonso-Prieto, 2011; see also Bobes et al., 2003; Dalrymple et al., 2011). Similarly, developmental prosopagnosics show typical activation across the “core” (posterior) regions of the face-processing system (OFA and FFA, e.g. Avidan et al., 2014). More recently, robust experimental techniques such as fast periodic visual stimulation (Liu-Shuang et al., 2016b) have been able to shed light on the important deficits in neural face-individuation of PS. Characterising the underlying computations of these neural and perceptual processes, however, remains a challenging task. First, describing neural computations is generally arduous due to signal-to-noise ratio (SNR) issues, which is even more concerning when recording brain activity from brain-lesion patients (Liu-Shuang et al., 2016a). Brain damage, for example, can significantly alter the flow of brain activity compared to typical observers, which can potentially deform event-related potential components (Alonso Prieto et al., 2011) and require more repetitions of conditions. Second, using solely temporal evidence is limited in itself to reveal brain computations as it only partially indicates the nature of the computations that are relied on by the brain (Lamme & Roelfsema, 2000; McDermott et al., 2002). Individuals relying on different neural computations in responses to faces, for example, could have identical activity at a given latency as indexed by univariate event-related potentials.

More explicit ways of revealing the nature of brain representations have recently gained traction with techniques associating functional and multivariate brain activity to computational models (Dwivedi et al. 2021; di Oleggio Castello et al. 2021; Popham et al. 2021; Kriegeskorte and Diedrichsen 2016; Doerig et al. 2022; Faghel-Soubeyrand et al. 2022). The aforementioned SNR concerns might explain why most work on prosopagnosia has relied on a limited set of stimuli conditions, block-designs, and univariate methods such as averaging of conditions and subtraction approaches. However, while significantly improving the statistical power of these studies, these approaches have prevented a thorough description of the brain computations underlying prosopagnosia (Friston et al., 2006). Investigating brain processing using condition-rich designs (Allen et al., 2022; Charest et al., 2014a; Kriegeskorte & Kievit, 2013; Naselaris et al., 2021), and promoting a broad description of underlying brain mechanisms by testing diverse models on a whole-brain basis (Dwivedi et al., 2021; Kriegeskorte & Diedrichsen, 2019; Popham et al., 2021) might provide a more comprehensive understanding of these processes.

Here, we take into account these temporal and computational aspects of brain processes by investigating prosopagnosia with fine-grained temporal recordings of brain activity (high-density electroencephalography; EEG), machine learning, and a proven multivariate method, i.e. Representational Similarity Analysis (Kriegeskorte, Mur, & Bandettini, 2008). We recorded the brain activity of PS and neurotypical controls in responses to images of various categories. Using multivariate pattern analysis (time-resolved “decoding”; Grootswagers et al., 2017), we probe PS’ neural evidence for face and non-face identity representations. Using RSA, we produce functional brain representations in a format that provides straightforward comparisons between individuals differing in neuroanatomical structure (Golarai et al., 2015; Popal et al., 2019). This enabled us to compare human brain representations with those of artificial models characterising different types of computations, i.e. deep neural networks of vision and semantic classification, thereby offering a window into the neural computations at play in patient PS’s deficits.

## Materials and Procedures

### Patient PS and neurotypical participants

A total of 20 participants were recruited for this study. The first group consisted of 19 neurotypicals individuals that included 15 healthy controls (9 female, Mage = 22.9 years old) as well as 4 aged-matched (3 female, Mage = 67.5). This sample size was chosen according to the effects described in previous mvpa studies (Carlson et al., 2013; Cichy et al., 2014; Faghel-Soubeyrand et al., 2022; Hebart et al., 2018), as well as previous studies on prosopagnosia (Gao et al., 2019; Humphreys et al., 2007; Liu-Shuang et al., 2016a; Richoz et al., 2015). Data from 10 of these participants (healthy controls 1-10) have been reported in a previous study (Faghel-Soubeyrand et al., 2022b). One participant from the aged-matched group (aged-matched #2) was rejected due to faulty EEG recordings and poor behavioural performance during the one-back task and CFMT+. This study was approved by the Ethics and Research Committee of the University of Birmingham, The University of Fribourg, and informed consent was obtained from all participants.

### PS’s case report

Patient PS was born in 1950 and is a *pure* case of acquired prosopagnosia. She was hit by the side mirror of a London’s bus in 1992 while crossing the road. This closed head injury led to major lesions in the left middle fusiform gyrus, where the left Fusiform Face Area (lFFA) is typically located, and in the right inferior occipital gyrus, which typically locates the right Occipital Face Area (rOFA; see Gao et al. 2019 for converging fMRI evidence). Both regions play a critical functional role within the face cortical network (Rossion, 2022a, 2022b). She also reported minor damages in the right middle temporal gyrus and left posterior cerebellum (for an exhaustive anatomical description and an illustration of her brain damages see, Sorger et al., 2007, Figures 2 and 3). Patient PS is a very well-documented and described case of acquired prosopagnosia. She has been extensively studied over the last 20 years, leading to impactful scientific contributions that significantly enriched the theoretical models on human face perception (Rossion, 2008, 2014; for a complete case report see, Rossion, 2022a, 2022b; Rossion et al., 2003b).

Patient PS recovered remarkably well from initially significant cognitive deficits with the support of medical treatment and neuropsychological rehabilitation. A couple of months after her injury she performed within the normal range at different non-visual tasks for which she was slightly impaired after the accident (e.g., calculation, short and long-term memory, visual imagery). Yet, her fine-grained visual discrimination abilities remained slower compared to controls, and she also presented reduced contrast sensitivity to high spatial frequency information (>22c/degree) and a profound prosopagnosia with massively impaired face recognition abilities (Rossion et al., 2003). The patient complains of a severe difficulty at recognizing faces, including the ones of close relatives (husband, children, friends), as well as her own face. PS can correctly categorise (and draw) faces as a unique visual object and discriminate faces from other non-face objects or scenes, even when the images are briefly presented (Schiltz et al., 2006). She shows no difficulty at object recognition, even for subordinate-level discriminations (Rossion et al., 2003; Schiltz et al., 2006). Patient PS is perfect at all tests from the Birmingham Object Recognition Battery (BORB – Riddoch & Humphreys, 2022) showing preserved processing of low-level aspects of visual information (i.e., matching of basic elementary features), intact object matching from different viewpoints, and normal performance for object naming (Rossion et al., 2003; Table 1). Her reading abilities are also well preserved although slightly slowed down, her visual acuity (0.8 bilaterally) is within the normal range, and her visual field almost intact apart from a small left paracentral scotoma. As reported by Rossion et al. (2003), she is highly impaired on the Benton Face Matching Test (BFRT - Benton & Van Allen, 1972) scoring 27/54 (percentile 1). She performs also poorly on the Warrington Recognition Memory Test (WRMT - Warrington & Shallice, 1984), scoring 18/25 (percentile 3) a performance that characterises her as impaired compared to controls. Over the years, patient PS developed strategies to infer a person’s identity by relying on external cues such as haircut, clothes, beard, glasses, gait, posture, or a person’s voice. Moreover, as revealed by the *Bubbles* response classification technique, patient PS uses suboptimal diagnostic information to recognise familiar faces, relying on the lower part of the face (i.e., the mouth region and external contours) instead of the most informative eye area (Caldara et al., 2005). A similar bias towards the mouth has been observed for the recognition of static facial expressions (Fiset et al., 2017) for which she is strongly impaired. Her ability to recognize the dynamic versions of the same facial expressions is nevertheless preserved (Richoz et al., 2015). Overall, PS is a very cooperative patient with extraordinarily preserved cognitive functions, sensory and motor skills and without any attentional deficits. She therefore represents an exemplary case to investigate the functional models of typical face processing.

### Behavioural tasks

#### One-back task

The stimuli used in the main experiment consisted of 49 images of faces, animals (e.g., giraffe, monkey, puppy), plants, objects (e.g., car, computer monitor, flower, banana), and scenes (e.g., city landscape, kitchen, bedroom). The 24 faces (13 identities, 8 males, and 8 neutral, 8 happy, 8 fearful expressions) were taken from the Radboud Face dataset (Langner et al., 2010). For further details on stimulus processing steps, see Faghel-Soubeyrand et al. (2022).

These stimuli were presented during a one-back task where we measured high-density electroencephalographic (EEG) activity (**Figure 1b,c**). Participants performed ∼3200 trials in two recording sessions, which were separated by at least one day and by a maximum of two weeks. Participants were asked to press a computer keyboard key on trials where the current image was identical to the previous one (repetitions occurring with a 0.1 probability). They were asked to respond as quickly and accurately as possible. Feedback about accuracy was given on each trial. A trial unravelled as follows: a white fixation point was presented on a grey background for 500 ms (with a jitter of ± 50 ms); followed by a stimulus presented on a grey background for 600 ms; and, finally, by a white fixation point on a grey background for 500 ms. Participants had a maximum of 1,100 ms following stimulus onset to respond. This interval, as well as the 200 ms preceding stimulus onset, constituted the epoch selected for our EEG analyses.

**Figure 1.**
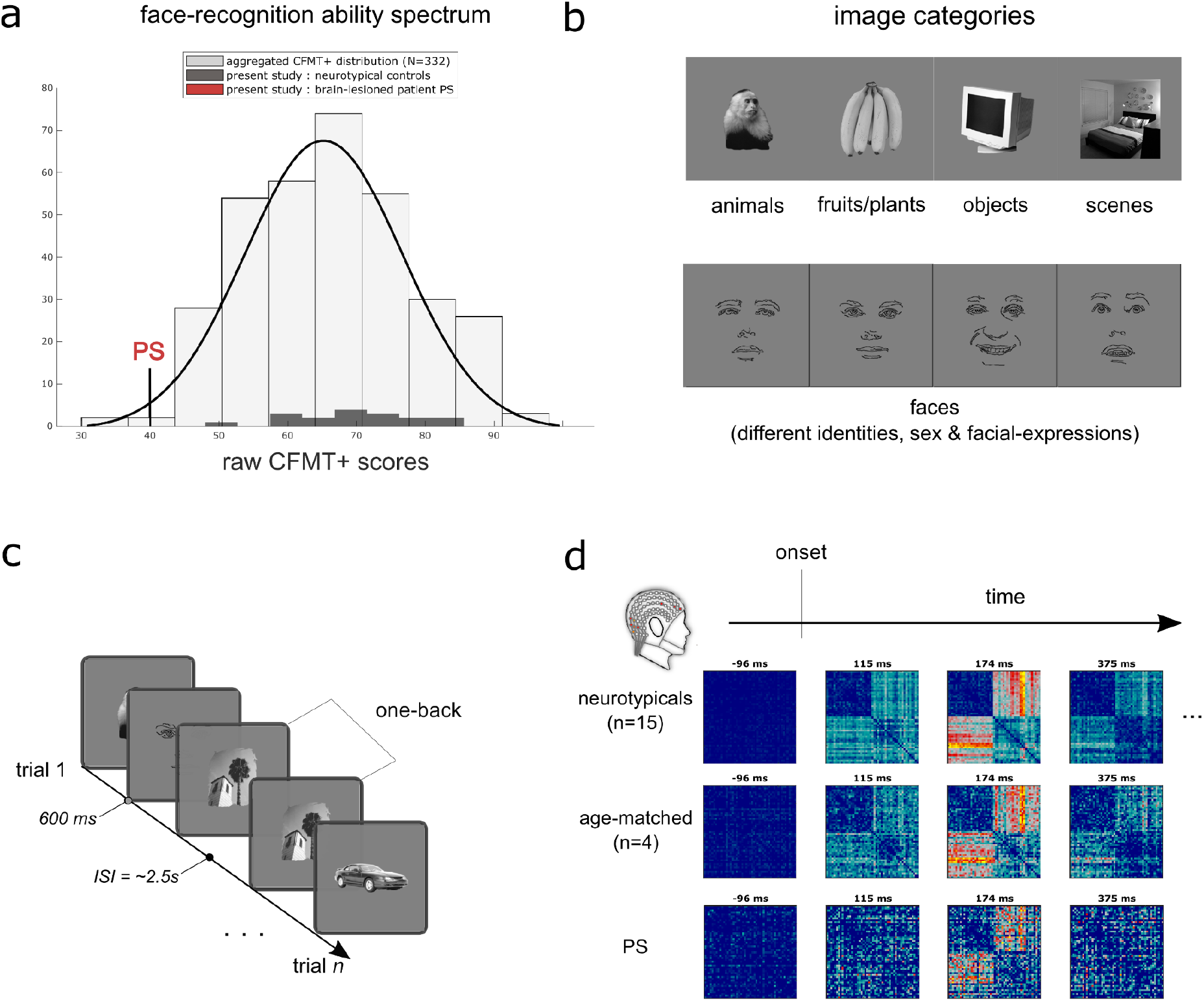
Overview of the experimental procedure. **a)** The histogram shows the Cambridge Face Memory Test long-form (CFMT+, Russell et al., 2009) scores of PS, typical recognisers (black bars), and an additional 332 neurotypical observers from three independent studies for comparison (Faghel-Soubeyrand et al., 2019; Fysh et al., 2020; Tardif et al., 2019). **b)** Participants engaged in a one-back task while their brain activity was recorded with high-density electroencephalography. The objects depicted in the stimuli belonged to various categories, such as faces, objects, and scenes. Note that the face drawings shown here are an anonymised substitute to the experimental face stimuli presented to our participants. **c)** Brain Representational dissimilarity matrices (RDM) were computed for PS and controls using cross-validated decoding performance between the EEG topographies from each pair of stimuli at every 4 ms time-point. Brain RDMs are shown for specific time points.

#### Cambridge Face Memory Test +

All participants were administered the CFMT long-form, or CFMT+ (Russell et al., 2009). In the CFMT+, participants are required to memorise a series of face identities, and to subsequently identify the newly learned faces among three faces. It includes a total of 102 trials of increasing difficulty. The duration of this test is about 15 minutes. EEG was not recorded while participants completed this test.

### EEG recording and preprocessing

High-density electroencephalographic data was continuously recorded at a sampling rate of 1024 Hz using a 128-channel BioSemi ActiveTwo headset (Biosemi B.V., Amsterdam, Netherlands). Electrodes’ impedance was kept below 20 µV. Data were collected at the University of Fribourg. Data was preprocessed using FieldTrip (Oostenveld et al., 2011) and in-house Matlab code: continuous raw signal was first re-referenced relative to A1 (Cz), filtered with a band-pass filter [.01-80 Hz], segmented into trial epochs from -200 ms to 1100 ms relative to image onset, and down-sampled at 256 Hz. These EEG recordings were completed during the one-back task only.

### Representational Similarity Analysis

We compared our participants’ brain representations to those from visual and caption deep neural networks using Representational Similarity Analysis (RSA; (Charest et al., 2014b; Kriegeskorte, Mur, & Bandettini, 2008; Kriegeskorte, Mur, Ruff, et al., 2008; Kriegeskorte & Kievit, 8/2013).

#### Brain Representational Dissimilarity Matrices

For every participant, we trained a Fisher linear discriminant (5-fold cross-validation, 5 repetitions; Treder, 2020) to distinguish pairs of stimuli from every 4-ms intervals of EEG response to these stimuli from -200 to 1100 ms after stimulus onset (Cichy & Oliva, 2020; Graumann et al., 2022). All 128 channels served as features in these classifiers. Cross-validated area under the curve (AUC) served as pairwise classification dissimilarity metric. By repeating this process for all possible pairs (1176 for our 49 stimuli), we obtained a representational dissimilarity matrix (RDM; see also **Supplementary figure 1a,b**). RDMs are shown for selected time points in **Figure 1d**.

#### Visual Convolutional Neural Networks RDM

We used a pre-trained AlexNet (Krizhevsky et al., 2012) as one model of the visual computations along the ventral stream (Güçlü & van Gerven, 2015). Our 49 stimuli were input to AlexNet. Layer-wise RDMs were constructed comparing the unit activation patterns for each pair of images using Pearson correlations (see also **Supplementary figure 1c** for a visualisation). This CNN process visual features of gradually higher complexity and abstraction along their layers (Güçlü & van Gerven, 2015), from low-level (i.e., orientation, edges in shallow layers) to high-level features (e.g., objects and object parts in deeper layers).

#### Caption-level Semantic RDM

To derive a model of an higher level of computations than purely visual processes, we first asked 5 new participants to provide a sentence caption describing each stimulus (e.g., “a city seen from the other side of the forest”) using the Meadows online platform (www.meadows-research.com). The sentence captions were fed as input in Google’s universal sentence encoder (GUSE; Cer et al., 2018) resulting in 512 dimensional sentence embeddings. GUSE has been trained to predict semantic textual similarity from human judgments, and its embeddings generalise to an array of other semantic judgement tasks (Cer et al., 2018). We then computed the dissimilarities (cosine distances) between the sentence embeddings across all pairs of captions, resulting in a caption-level semantic RDM for each participant. The average RDM was used for further analyses (see also **Supplementary figure 1c** for a visualisation).

### Decoding analyses

#### Face-identity decoding

We trained multiclass linear discriminant classifiers to predict 8 face identities (5-fold cross-validation, 5 repetitions; Grootswagers et al., 2017; Treder, 2020), using all 128 channels single-trial EEG data as features. A sliding-average with a window of 39 ms was applied to EEG traces prior to training. Separate classifiers were trained on the resulting successive 4-ms EEG time intervals. The classifiers were trained on trials of EEG activity of participants viewing face-identities varying in facial-expressions (fear & joy; ∼525 observations per session per participant), similarly to previous studies of face-identity information using EEG signal (Nemrodov et al., 2016). The time courses of these decoders’ performance were averaged across sessions. Recall (defined as : *true positives* / [*true positives* + *false negatives]*) was used to assess decoding performance.

#### Non-face identity decoding

PS’s impairments for individuation of visual objects are restrained to faces (Rossion, 2018, 2022a). As a standard of comparison to the neural face-identification decoders, we trained multiclass linear discriminant classifiers decoders to predict 8 within-category identities from *non-face* categories (i.e. objects, scenes, and animals). These classifiers were trained with identical parameters to the face-identity classifiers. Separate classifiers were trained for each category (e.g. a decoder for 8 identities within the animal category, another for 8 identities within the object category, another for 8 identities within the scene category; ∼260 observations per session per participant). The time courses of these decoders’ performance were averaged across all three categories and sessions. Recall was again used to assess decoding performance.

### Comparison of brain representations with computational models

We compared our participants’ brain RDMs to those from the vision (**Figure 3a**) and caption-level description (**Figure 3b**) models described in the previous section using Spearman correlations. Partial correlations were used where mentioned.

### Reliability of brain representations across recording days

We computed the reliability of brain representations in a similar way as in Charest et al., 2014 (Charest et al., 2014a). For each participant, two RDMs were computed from two separate recording days, and compared using Spearman correlation in a time-resolved manner. Significance was assessed using permutation testing. Specifically, we created, for each participant and at each 4 ms step, a null distribution of 1000 brain-to-brain correlations using an RDM in which rows and columns indices were randomly shuffled.

### Group comparison and inferential statistics

All contrasts between PS and neurotypical controls were computed using Crawford-Howell modified t-tests for case-controls comparisons (Crawford & Garthwaite, 2012; Crawford & Howell, 1998). All time-resolved contrasts were computed from 0 to 1.1 s after image-onset. Spearman correlations were used for correlations within the control group, i.e. correlation analyses with behavioural performance.

Permutation testing was used throughout the paper to assess significance of EEG-RDM to computational model latencies. We created, for each participant and at each 4 ms step, a null distribution of 1000 brain-model correlations using model RDMs in which rows and columns indices were randomly shuffled (Kriegeskorte, Mur, & Bandettini, 2008).

Permutation testing was also used to assess significance of identity decoding latencies. We created, for each participant, session, and at each 4 ms step, a null distribution of 500 decoding performances using identity labels which indices were randomly shuffled. The average of these distributions across sessions and participants were used to assess significance of the time-resolved decoding performance shown in **figure 3a**.

## Results

### One-back task

Accuracies did not differ between aged-matched and young controls sub-groups either for face stimuli (t(16) = -0.3099, *p* = .761; t(16) = -0.9607, p =.3510) or non-face stimuli (t(16) = 1.2925, *p* = .215; t(16) = -.09704, p = .3463). Therefore, their data have been aggregated into a single neurotypical control group. PS differed significantly from controls on a face vs. non-face performance score, computed as the first PCA component of face vs. non-face accuracies and response times (t(17) = -7.7157, p = 1.6053×10-6).

### Cambridge Face Memory test long-form (CFMT+)

Within the control participants, CFMT+ scores did not differ between aged-matched and young controls sub-groups (t(16) = -0.8058, *p* = .4322). Their data have been aggregated into a single control group. PS significantly differed from controls on this standard face identification ability score (t(17)=-2.7623, *p* = 0.0133).

To assess the face-specific performance of PS and controls in a single individual score across all behavioural measures, we combined performance in the one-back task (accuracies and response times of face and non-face trials) and face-memory performance (CFMT+) using Principal Component Analysis (PCA). Specifically, face-specific performance in the one-back tasks was computed as a face vs. non-face performance score ([face - nonface]./[face + nonface]) separately for accuracy and RTs, for each participant. We used PCA to extract projections explaining variance across these two variables as well as the CFMT+ (Calder et al., 2001; Calder & Young, 2005). The first component, which explained 65.76% of the variance in performance across participants, is henceforth referred to as the face-specific performance score. PS significantly differed from neurotypical controls on this score (t(17) = -7.1966, *p* = 1.0691e-06; see **Figure 2a**), indicating typical face-specific behavioural deficits in this patient.

**Figure 2.**
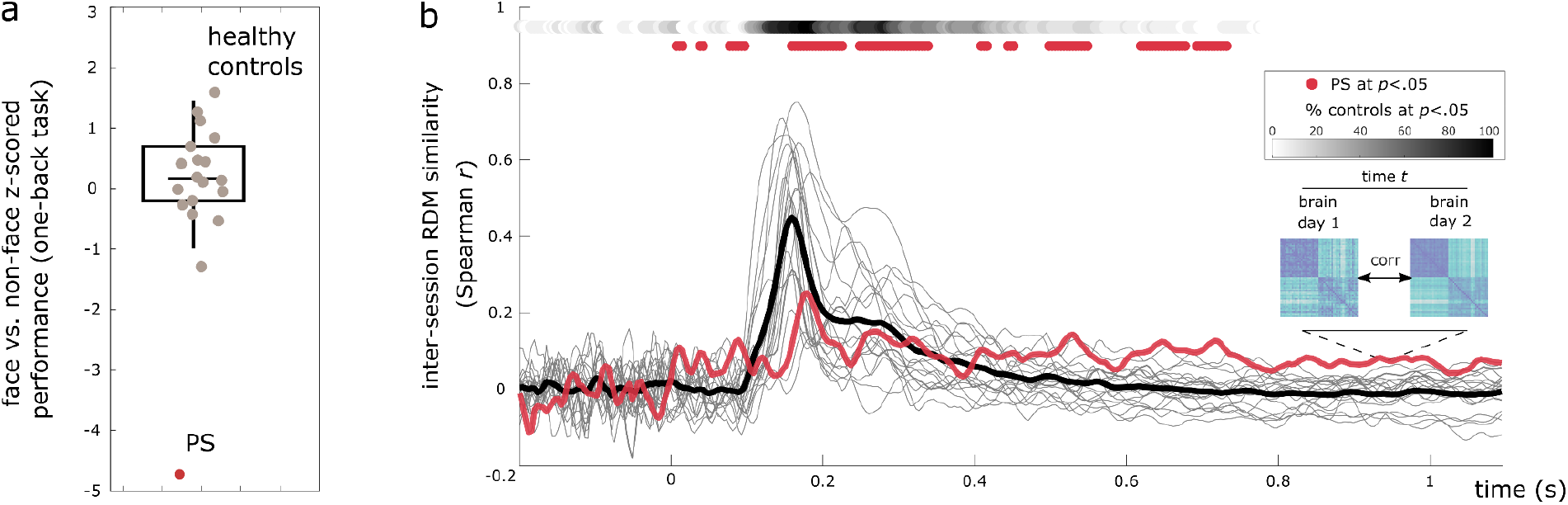
Behavioural performance and reliability of brain representations across time. **a)** A behavioural face-specific performance score was computed for all participants and constrasted between PS and controls, indicating strong and archetypical face-sensitive deficits (*p*<.0001). **(b)** Brain Representational Dissimilarity Matrices (RDMs) computed on different recording days were cross-correlated within participants for patient PS (red line) and neurotypical participants (grey lines indicate individuals participants, black line indicate control-averaged). PS brain RDMs showed significant inter-session reliability across most time windows (*p*<.05, permutations). For comparison, the percentage of control participants with significant cross-session correlations is shown on top for every time point, with darker points indicating higher percentage of neurotypicals with reliable brain RDMs. PS had similar reliability coefficients compared to controls.

### Reliability of neural dynamics measured with RSA

Measuring brain dynamics of brain-lesioned patients can be arduous using raw electrophysiological topographies (Alonso Prieto et al., 2011). We assessed whether we could measure reliable brain representations of brain-lesioned PS by computing inter-session reliability of brain Representational Dissimilarity Matrices (RDMs). We computed the correlation between EEG RDMs computed from recording day 1 and recording day 2 at every 4 ms steps. The timecourse of these correlations, shown in figure 2a, indicates significant (*ps*<.05, permutation testing) reliability of brain representations of both neurotypicals and PS across most time points after image onset, peaking in the N170 window (160 ms and 180 ms for controls and PS, respectively; r_peak_ctrls_ = .4316; r_peak_PS_ = .2638). PS showed surprisingly high SNR across sessions, her correlation time course being indistinguishable from the neurotypicals’ reliability scores across time points (i.e. no significant contrasts).

### Impaired neural decoding of face-identity

We first asked whether we could capture PS’s face identification deficits at the neural level (Gao et al., 2019; Liu-Shuang et al., 2016). For each participant, we decoded face-identity from brain activity by training a multiclass linear discriminant classifier from whole-brain high-density EEG patterns at each 4 ms steps from face onset (see **Figure 3a**). In neurotypical controls, this resulted in weak but above chance decoding of face identity across time (from 138 ms after face onset, peaking at around 180 ms; *ps*<.05, permutations; see **Figure 3a**), as can be expected from this difficult classification task (Kriegeskorte et al. 2007; Tsantani et al. 2021; Muukkonen et al. 2020; Dobs et al. 2019). PS’s neural face-identity decoding was continuously at chance level (no significant time points after face onset; *ps* >.05, permutations). Time-resolved contrasts with controls confirmed the reduced neural identity decoding in PS around 200 ms (Howell-Crawford t-tests; *ps*<.05). Neural decoding of *non-face* identities, on the other hand, appeared entirely spared in PS. Non-face identity classifiers resulted in above chance performance in both controls (from 84 ms to 620 ms after non-face image onset, peaking at around 140 ms; *ps*<.05, permutations) *and* PS (from 80 ms to 290 ms, peaking at around 228 ms; *ps*<.05, permutations). PS’s neural individuation for non-face objects was also within the normal range of controls across all time points after face onset (no significant contrasts PS < controls; see **Figure3b**). Peak individual neural decoding evidence for face identities (**Figure 3c**) and non-face identities (**Figure 3d**) across participants demonstrated the same face-specific pattern, with PS positioned at the low end of the spectrum of the neural face individuation scores (*t*(18) =-2.48, *p*<.05), but showing her within typical scores on the neural *non-face* individuation evidence (*t*(18) = -0.73, *p* =.47).

We further tested the behavioural relevance of this decoded face identity evidence to the general population. We correlated the magnitude of neural decoding across participants (i.e. the neural individuation score) with behavioural face-identification abilities, as measured by the CFMT+ within our sample. Notably, this was done *with individuals from the control group only*. This showed that, indeed, neural face-individuation score correlated positively with face recognition abilities across neurotypicals (peak-*r* = 0.63, peak-*r*_time_ *=* 330 ms; *ps*<.05, ∼174-702 ms; **Figure 3c**; *p*<.05, permutations). Identical brain-behaviour correlation analyses using *non-face* neural individuation scores resulted in much reduced correlations with the CFMT+, with mid and late correlation windows disappearing and the only significant positive correlations appearing in a short window around 100 ms after non-face objects onset (**Figure 3d;** *p*<.05, permutations). Both the contrasts between PS vs. controls and these interindividual results were replicated using the brain RDMs of our participants and identity model RDMs (Dobs et al., 2019; see **Supplementary material**).

**Figure 3.**
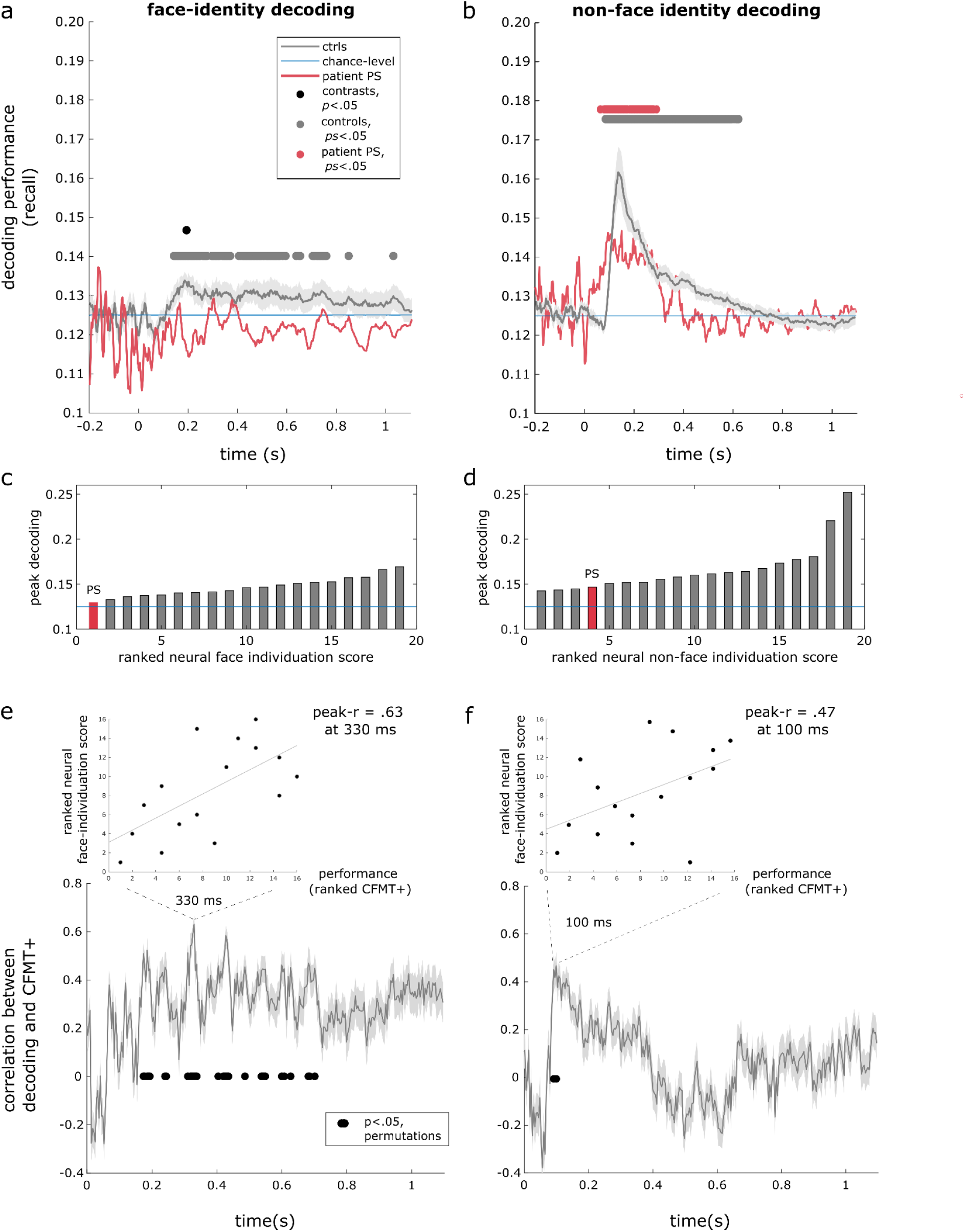
Neural decoding of face and non-face identities and association with individual face recognition abilities. Multivariate classifiers were trained to predict 8 face identities **(a)** as well as 8 identities from non-face categories **(b)** from time-resolved EEG patterns. **a)** Neural face-identity decoding performance (recall, 5-fold, 5-repetition cross-validation; Treder, 2020) is shown for PS (red line) and the control group (black line, representing control-average decoding time course; grey dots show significant identity decoding at *p*<.05, permutations). PS had overall below chance decoding for the face-identity decoding across time and showed significantly lower identity decoding compared to controls around 200 ms after face onset (black dots indicate significant contrasts PS < controls at *p* <.05). Shaded error bar represents standard error within the control group. **b)** PS showed typical neural decoding of non-face identities across all time points, with no significant contrasts with controls. Peak individual neural decoding evidence for face identities **(c)** and non-face identities **(d)** have been ranked across participants, and demonstrated similar effects with PS at the lower end of the spectrum of the neural face individuation scores, but was within typical scores on the neural non-face individuation. **e)** Spearman correlation between individual neural decoding evidence and face recognition abilities (i.e. the CFMT+ scores across neurotypical participants only) was computed at each 4 ms step. Correlation time course and shaded error bar were computed using a jackknife procedure. Significant brain-behaviour correlations (permutation testing at *p*<.05) were found in several time points for the face-identity decoding (between 174 ms and 702 ms, peaking at 330 ms). **f)** In contrast, brain-behaviour correlation with non-face identity decoding and CFMT+ scores were restrained to a small early window peaking at 100 ms (92-100 ms). Each panel shows a scatter plot of CFMT+ and neural decoding computed at peak correlation with corresponding least-square regression line.

Thus, we were able to capture the poor neural identity representations of PS across time using multivariate EEG signal, showing her reduced neural distance between face identities around 200 ms. This impairment appeared specific to face individuation, and the magnitude of this representational distance, particularly around 330 ms, was predictive of neurotypical face identification abilities.

### Similarity with visual and semantic computational models

Next, we characterised the specific neural computations underlying these deficits. We assessed visual brain computations in PS and neurotypical controls by comparing their brain RDMs to those of convolutional neural networks (CNNs) trained to categorise objects (Güçlü & van Gerven, 2015; Krizhevsky et al., 2012; Simonyan & Zisserman, 2014). The visual model RDMs were produced for all 8 layers of the CNN (see **Supplementary figure 1**). Controls showed significant brain-RDM correlations with the sixth layer of the visual CNN around 98 ms (CNN layer 6, *p*<.05, permutations), continuing as late as around 500 ms after image onset (see **Figure 4a;** CNN layer 3 : 59 - 746 ms; CNN layer 1 : 100 - 242 ms). PS showed significant correlations with the CNN layer 6 in a more restrained window between 153 and 204 ms (layer 1: ns; layer 3: 211 - 215 ms). Direct contrasts of PS’ correlation time courses with those of controls indicated reduced correlations with RDMs across the visual CNN (i.e., at layers 1, 3, 6 & 7; **Figure 4a**, *p*<.05). These contrasts peaked at layer 6, which represent a higher proportion of high-level visual features (e.g., whole objects and object parts, Güçlü & van Gerven, 2015; Long et al., 2018).

**Figure 4.**
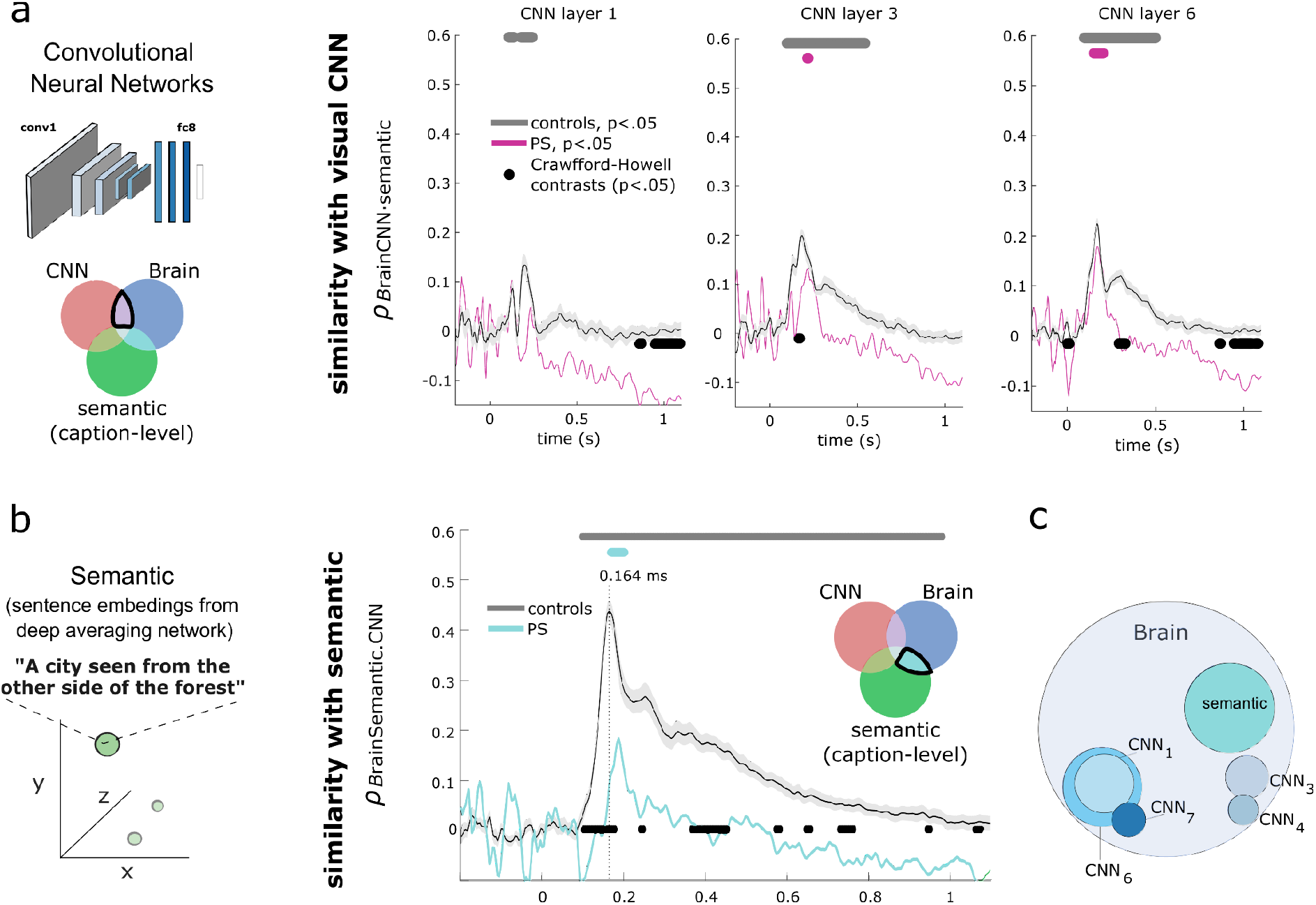
Comparison of brain representations with those of artificial neural networks of visual and semantic processing. **a**) Partial correlation between brain RDMs and AlexNet RDMs (removing shared correlation between brain and semantic model) is shown for PS (coloured curve) and controls (grey curve). Each column shows different layer RDMs in ascending order from left to right. We found overall lower similarity of visual computations within the brain of PS compared to controls (black dots indicate significant contrasts, Howell-Crawford modified t-tests, *p*<.05), with differences peaking in higher-level CNN layer 6. Similar results were observed when comparing brains and CNN models without removing the shared information between brains and the semantic (caption-level) model (see supplementary figure 3). **b**) Partial correlation with RDMs of the semantic model (excluding shared information between brain and AlexNet) was significantly lower in the brain of PS compared to controls (cyan curve; black dots indicates significant contrasts, *p*<.05). The shaded areas of all curves represent the standard error for the controls. **c**) Proportion of all time points after EEG onset showing significant contrasts (PS < controls, *p*<.05, uncorrected) is shown on a Venn diagram as a summary of PS’s impaired brain computations for visual computations (blue colours, where deeper blue represents deeper CNN layers) and semantic computations (cyan). The total surface area for the “brain” section represents all possible time points after onset (0-1100 ms in 4 ms steps), while the surface of each categories represents a portion of these time points. Overlap indicates significance over the same time points.

To reveal whether even higher-level semantic computations (Barton et al., 2009; Schweinberger & Neumann, 2016) could be affected in the brain of PS, we used a deep averaging network (Google Universal Sentence Encoder, GUSE; Cer et al., 2018) to transform human-derived captions of our stimuli (e.g. “ a city seen from the other side of the forest “) into embeddings (points in a caption space). Then, we compared the RDMs computed from this semantic model to the brain RDMs of PS and control participants. Controls showed significant correlations with semantic computations from around 98 ms, continuing as late as 981 ms after image onset, and peaking at 164 ms (peak-*r*_ctrls_ = .45; *p*<.05, permutations, **Figure 4b**). Again, PS showed significant correlations with this model in a more restrained window from around 176 to 200 ms, and peaking ∼20 ms later than controls at 188 ms (peak-*r*_PS_ = .173; *p*<.05, permutations, **Figure 4a**). Direct contrasts confirmed the reduced correlations with these semantic computations in the brain of PS compared to controls (**Figure 4b**, *p*<.05). Note that this comparison, as well as the one with the visual model, excluded the information shared between the semantic and visual models. Similar results were observed when comparing brains and the visual and semantic model without removing this shared information between brain and visual/semantic CNN (see **Supplementary figure 3 & Supplementary figure 4**).

A summary of significant contrasts with all computational models (**Figure 4c**) indicates that PS’s brain processing stream exhibited impairments peaking in higher-level visual (CNN layer 6) and semantic (caption-level) representations.

### Relationship with electrophysiological brain components

We further specified the time at which visual and semantic brain computations differed in PS by producing brain-to-model similarities in time windows corresponding to well-known Event-Related Potential (ERP) components indexing early, mid, and late neural processing, i.e. the P100 (Luck et al., 1990), N170 (Bentin et al., 1996) and N400 components (Kutas & Federmeier, 2000), respectively (**Figure 5a**). Within neurotypicals, we found EEG representations peaking in similarity with the visual CNN at mid-layers (fourth and fifth; Jiahui et al., 2022) around mid-level temporal windows. Similarity with semantic computations also peaked around mid-latencies. EEG correlations with the semantic model, however, far surpassed those with the visual CNN at mid and late latencies (paired t-tests comparing brain-semantic vs. brain-visual correlation, *ps* <.05).

**Figure 5.**
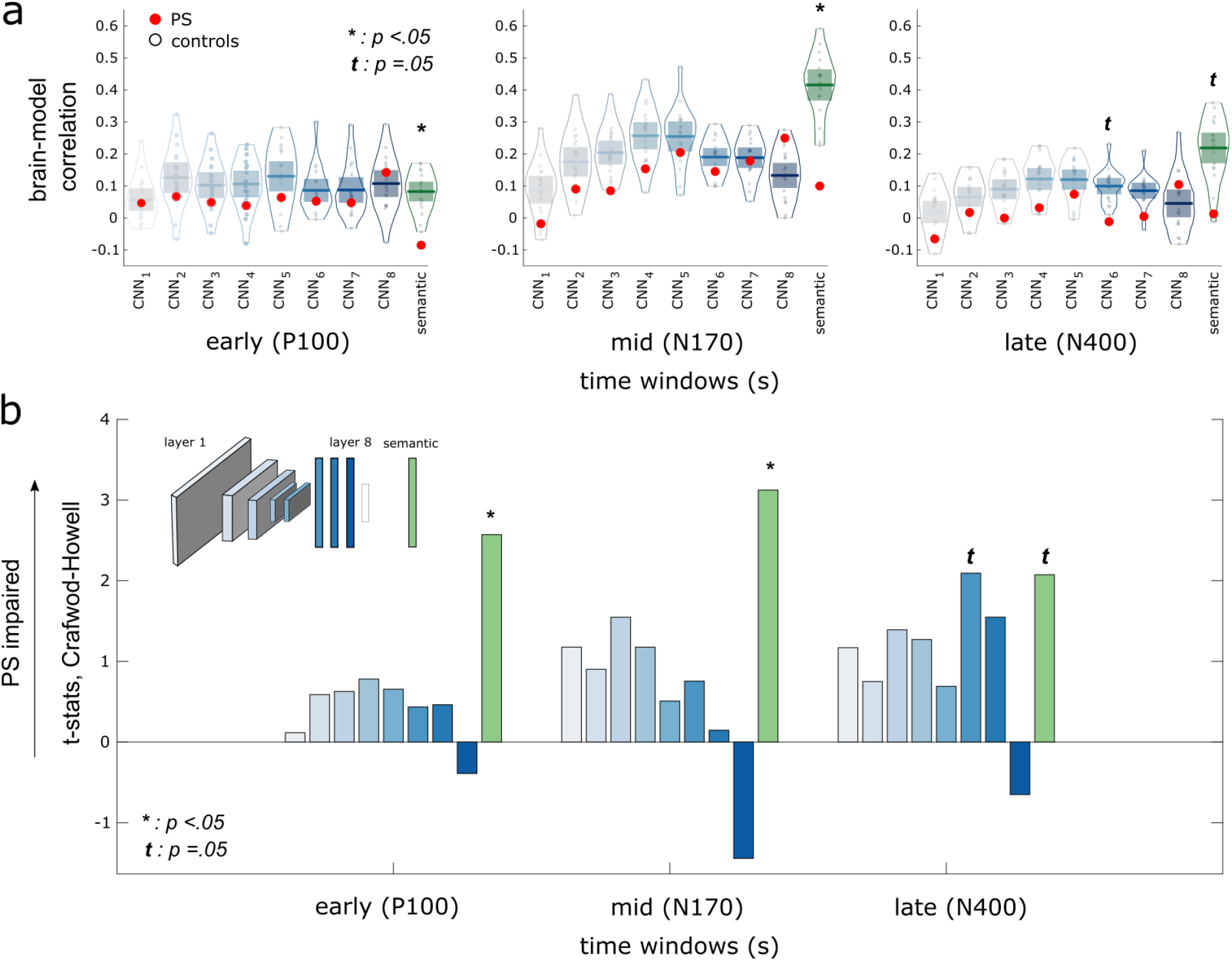
Comparison of visual and semantic computations along early, mid and late brain processing steps. **a)** Correlations of brain RDMs to visual model (blue plots, with deeper blue corresponding to deeper CNN layers) and semantic model (green plot) was assessed in PS and controls for brain RDMs computed in temporal windows corresponding to the P100 (80-130 ms), N170 (130-230 ms) and N400 (250-500 ms) ERP components. **(b)** Crawford-Howell t-statistics comparing controls vs. PS on these correlations are shown for the visual CNN (blue bars, with deeper blue corresponding to deeper CNN layers) and the semantic model (green bars; significant comparisons are indicated with an asterisk, marginally significant comparisons with the letter *t*).

In contrast to this tendency, PS showed generally lower similarity to semantic computations across all time windows. Direct contrasts of brain-model similarity between controls and PS are shown in **Figure 5b**, with higher values indicating stronger impairments in PS in visual/semantic computations. We observed reduced semantic brain computations for early and mid-processing windows around the P100 and N170 components (Bentin et al., 1996; Clark et al., 1994), respectively, as well as a marginally significant effect in late windows around the N400 component (*p* = .0536; Kutas & Federmeier, 2000). Contrasts of similarity with the visual CNN layers were overall less pronounced after averaging RDMs in these ERP windows. Similarity to the CNN layer 6 showed the only marginally significant difference between PS and controls around the late processing window corresponding to the N400 (*p* = .0512). More fine-grained temporal contrasts using time windows of 60 ms steps (see **Supplementary figure 5**) confirmed these results.

As stated earlier, a predominant assumption in cognitive-neuroscience is that temporally early brain signal refers to low-level computations while later brain signal refers to higher-level computations (e.g. Wiese et al. 2019; DiCarlo et al. 2012). To assess whether such progression was present in the neural time course of our participants, we computed all pairwise correlations between brain RDMs, CNN-layers RDMs, and semantic RDMs, resulting in a second-level representational similarity matrix (RSM) of 12 × 12 correlation values (**Figure 6a**). Comparing all RDMs in this manner is similar to the analyses described in the previous section, but allows two additional insights. First, it reveals how similar brain representations computed from different temporal windows are with one another (King & Dehaene, 2014). In our setting, this showed that neural representations during the N170 are more similar to those during the N400 compared to earlier (P100) computations within controls. Second, this analysis reveals how similar the representations of computational models are along their putative hierarchical computational stream (i.e. layers 1-8 & semantic processes; Güçlü & van Gerven, 2015). This showed, reassuringly, that the representations of the CNN used here are also generally more similar in layers closer apart within the CNN’s architecture (e.g. layer 6 and 7 are more similar to one another than layer 1 and 7). Finally, both of these levels of analysis *and their interaction* can be summarised by computing the optimal two-dimensional solution of this multidimensional space with multidimensional scaling (MDS). This visual summary is shown in **Figure 6b**. Because it produces 2D coordinates in a (representational) space as it unfolds across time, producing a path or progression of brain representations, it will be referred to as representational trajectory. In both PS and controls, this analysis confirmed a trajectory from early brain representations, relatively more similar to early layers of the visual CNN, to mid and late brain representations, relatively more similar to the deeper layer of the CNN and semantic model. Some differences, however, are noticeable. First, these second-level RSMs indicated the reduced magnitude of similarity to the models in PS outlined in the previous section, specifically to those of higher-level semantic computations. More importantly, these representational trajectories revealed that PS’ late neural code is both i) more similar to her early (brain) representations around the P100, and ii) more similar to low-level (visual) computations of the CNN. These results indicate that PS shows relatively less transformations in neural computations from early to late stages of her visual processing stream (Cichy & Oliva, 2020).

**Figure 6.**
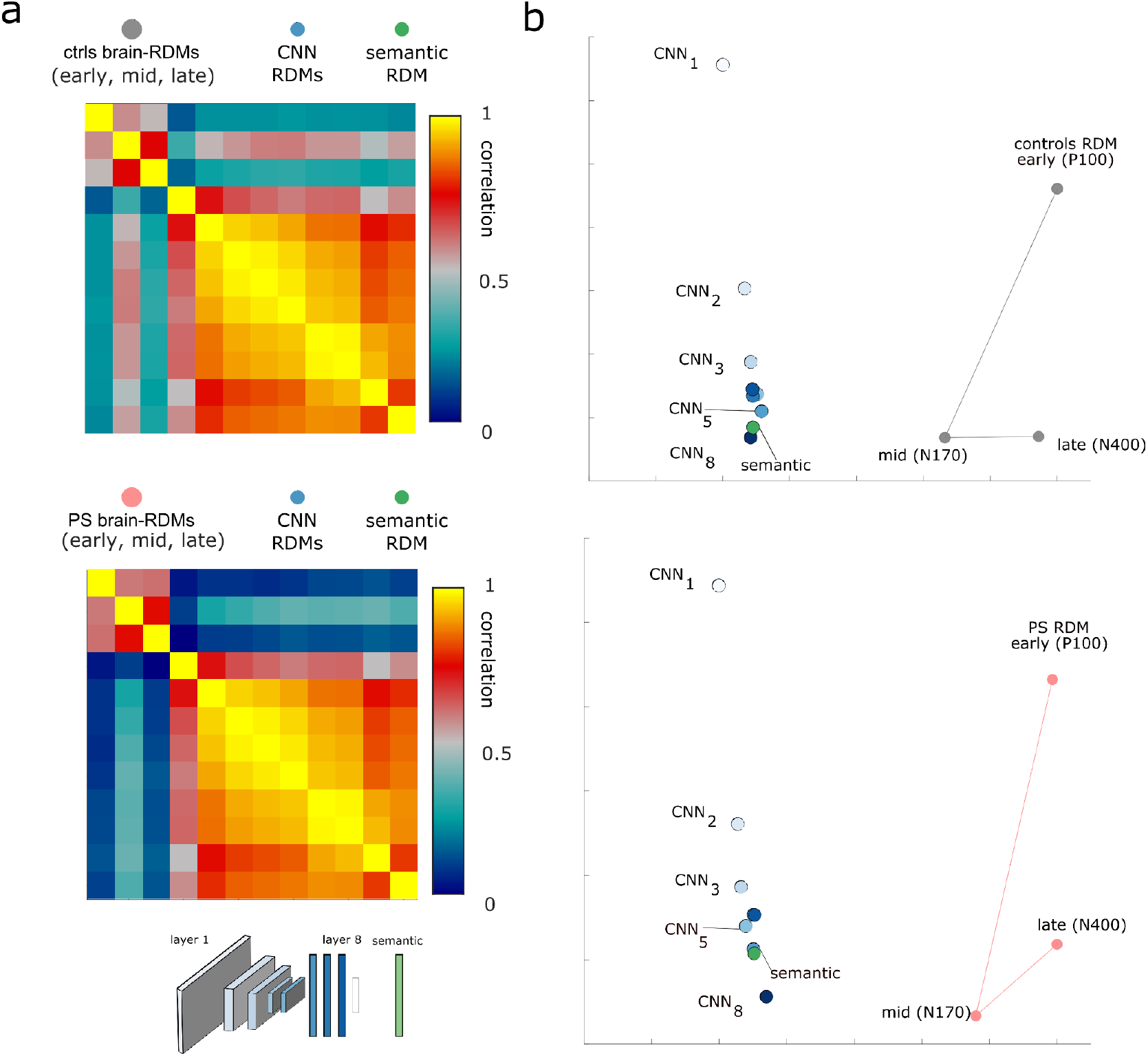
Representational trajectories of brain and computation models. **a)** All pairwise correlations across brain and computational models’ RDMs resulted in second-order similarity matrices (RSMs) for controls (upper panel) and PS (lower panel). **(b)** A summary of group RSMs was obtained by computing the 2D solution of these RSMs using MDS. Points corresponding to brain representations of controls (upper panel) and PS (lower panel) were linked with grey and red lines, respectively.

## Discussion

Here, we characterised the neural computations of a well-studied single-case of acquired pure prosopagnosia, patient PS, using whole brain multivariate signal and computational models of vision and caption-level semantics. We were able to capture the impaired functional neural signatures of face individuation — as well as the absence of non-face individuation deficits — in the brain of PS. Most importantly, we show that PS’ inability to identify faces is associated with reduced high-level visual computations, including a significant reduction of even higher-level semantic brain computations. This reduction in semantic computations was present as soon as the P100, peaked around the well-known N170 component, and persisted later around the N400 window. Analyses of brain dynamics revealed that the neural code of patient PS appears to show reduced transformations in neural computations across time, with fewer changes from early to late brain representations.

Impairment for face individuation is the archetypal deficit of patient PS and prosopagnosia in general (Bodamer, 1947), but has been arduous to reveal solely from brain activity (Anzellotti et al. 2014; Alonso Prieto et al. 2011). Using a data-driven whole-brain approach and high-density EEG, we extend recent work having been able to reveal such deficits from the neural responses of PS with fast periodic visual stimulation (FPVS; Liu-Shuang et al., 2016) by describing the temporal unfolding of this deficit under normal passive viewing conditions. We found reduced neural evidence for face identity around the N170 window (Bentin et al., 1996) in PS, and, conjointly, revealed normal *non-face* individuation in this patient (Dricot et al., 2008; Liu-Shuang et al., 2016). We further discovered that variations in *typical* face recognition abilities, as indexed by gold-standard behavioural tests in control participants (Russell et al., 2009), is associated with this neural decoding of face identities from around the N170 onward, peaking around 350 ms in the N400 window (Johnston et al., 2016; Schweinberger & Neumann, 2016; Wiese et al., 2019). Again, individual neural decoding of non-face individuation did not correlate with neurotypical face-recognition abilities in these time windows but was rather only associated with early neural activity around 100 ms after face onset.

Crucially, by associating electrophysiological signal and computational models, we found that the underlying neural computations of PS differed most with respect to the higher-level visual and semantic computations of deep neural networks models (DNNs). The late layers of visual DNNs have been previously linked to processing in human infero-temporal cortex (hIT; Güçlü & van Gerven, 2015; Jiahui et al., 2022.; Khaligh-Razavi & Kriegeskorte, 2014), peaking in the FFA (Khaligh-Razavi & Kriegeskorte, 2014), and functionally to higher-level visual feature representations such as parts of objects, whole objects and viewpoint invariant representations (Güçlü & van Gerven, 2015). These observations are consistent with the impaired whole face (Ramon et al., 2016) and feature representations (Caldara et al., 2005; Fiset et al., 2017) previously described in patient PS. Associations between brain activity and higher-level semantic computations have only been attempted much more recently (Dwivedi et al., 2021; Faghel-Soubeyrand et al., 2022; Popham et al., 2021). The computations revealed here are arguably closer to visuo*-*semantic representations, which have been shown to be represented both within the visual ventral stream (including the FFA and OFA; Doerig, Kietzmann, et al., 2022; Popham et al., 2021) and outside (Doerig et al. 2022). Using a model of caption-level semantics (Cer et al., 2018; Faghel-Soubeyrand et al., 2022; Doerig et al. 2022), we show that PS’s representational geometry displays significant — but greatly reduced — correlation with these semantic computations compared to controls. These findings demonstrate a clear link between semantic brain computations and important changes in the ability to recognise faces (Bruce & Young, 1986; Duchaine & Yovel, 2015).

Together with similar computational characterisation of brain representations in individuals with “super-recognition” of faces (Faghel-Soubeyrand et al., 2022; Russell et al., 2009), our results suggest *some* gradient of neural computations from the low-end to high-end of face-recognition abilities. Indeed, super-recognisers in Faghel-Soubeyrand et al., (2022) showed *enhanced* similarity with mid-level visual and semantic computations around the N170 and P600 windows, respectively. Here, while PS showed reduced similarity with visual and semantic computations, obvious differences are also apparent. PS’s neural computations, particularly semantic computations, appear to be impacted much sooner and on a larger extent than those of super-recognisers. Whereas semantic brain computations were enhanced around the P600 in super-recognisers, here PS displayed reduced similarity as early as the P100, continuing throughout the N170 and N400 windows. In fact, PS showed no significant correlation with either the visual or semantic-level computations at any point during late processing windows (i.e. after ∼300 ms). These observations suggest that PS’s deficits in neural computations start relatively early along the typical processing stream. Some differences, however, are to be expected given the important lesions to the face processing network (i.e. including the right-OFA and left-FFA) of this patient. Indeed, the OFA is not only involved in the extraction of featural information about faces (Pitcher et al., 2007), but has also been causally linked to higher-level identity (Ambrus et al., 2017; Jonas et al., 2014) and semantic information (Eick et al., 2020) processing. Both these associations and the fact that the Anterior Temporal Lobe (ATL) — an important hub for the processing of semantic information — is preserved in PS (Rossion 2022; Gao et al. 2019) points to a similar involvement of the OFA/FFA complex in the semantic neural computations revealed here. Future work using brain imaging with higher spatial resolution will be required to address these questions.

While the present study offers insights on the neural dynamics and computations of prosopagnosia, it is also limited in a number of ways. For example, even though we recorded a sizable data set of high-density EEG comprising numerous trials and stimulus conditions, signal-to-noise ratio was still small when probing specific neural processes. Characterising face identity representations across time is a notoriously difficult task (Anzellotti et al., 2014; Dobs et al., 2019) that could have benefited from recordings of even higher density imaging (M/EEG in Dobs et al., 2019; Vida et al., 2017; fMRI in Nestor et al., 2011), as well as from the use of experimental tasks recruiting more directly this process (Nemrodov et al., 2016). Another related and potentially fruitful approach to reveal brain computations and representational geometry in prosopagnosia would be to produce EEG-fMRI “fusion” (Cichy et al., 2014; Cichy & Oliva, 2020). Indeed, the spatio-temporal description procured by linking EEG *and* fMRI data with RSA would be ideal to test hypotheses regarding transformations in neural code such as the one hinted in our findings. Employing this technique on PS’s EEG and additional fMRI data could reveal the putative hurdles in the transformation of neural code across the different lesions of PS’s, and thereby attribute causal links between OFA/FFA and impairments in brain computations. Finally, comparison of brain representations with computational models in general also has certain limits (Doerig, Sommers, et al., 2022). Recent advances in developing more interpretable (Schyns et al., 2022; Soulos & Isik, 2020) and ecological models (Kietzmann et al., 2019; Mehrer et al., 2021) of human cognition will be helpful to address them.

Notwithsanding these limitations, this work offers, to our knowledge, the first description of the fine-grained temporal processes combined with a state-of-the-art computational characterisation of the brain representations in prosopagnosia. Understanding the very nature of defective perceptual representations offers new routes for patient rehabilitation. Indeed, we believe that the recent technological advances that permitted us to reveal these findings offer new promising ways to understand, and could perhaps even help diagnose the fine-grained deficits in perception and cognition in diverse clinical populations.

## Acknowledgements

We sincerely thank PS for her precious contribution and participation in this study.

## Author contributions

(CRediT standardised author statement)

**S.F-S**. : conceptualisation, methodology, software, formal analysis, investigation, data curation, writing - original draft, visualisation, supervision, project administration, funding acquisition. **A-R.R**. : project administration, investigation, writing - editing and reviewing. **J.W**. : funding acquisition. **D.W**. : Investigation **R.C**. : resources, project administration, writing - editing and reviewing. **F.G**. : supervision, writing - editing and reviewing. **I.C**. : supervision, methodology, resources, formal analysis, writing - editing and reviewing, project administration, funding acquisition.

## Data availability

Data are available from the corresponding authors upon request.

## Code availability

The MATLAB and Python codes used in this study will be available upon request.

## Supplementary material

**Supplementary figure 1.**
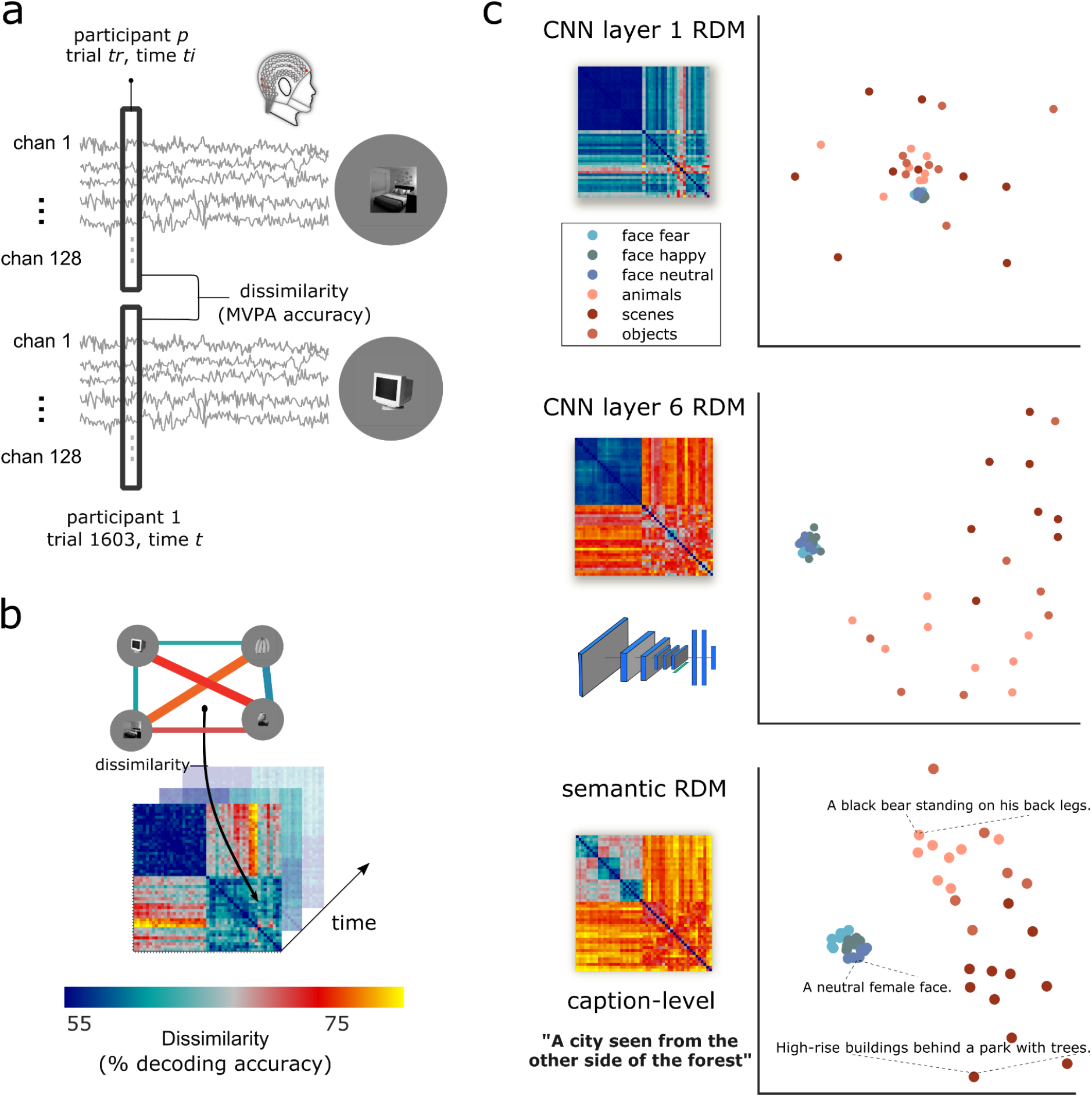
Creation of high-density EEG RDMs and visualisation of computational models RDMs. **a)** The cross-validated dissimilarities between raw topographies from all pairs of stimuli presented to our participants were used to produce brain EEG RDMs **(b)**. RDMs of example CNN layers 1 and 6 as well as the RDM for the caption-level semantic model are also presented in **(c)**, with their corresponding 2D visualisation using Multidimensional Scaling (MDS).

### Neural dynamics for key categorical distinctions using RSA

We derived neural distances between face-identities by correlating the brain RDMs of our participants with a categorical face identity model RDM (where ones indicated multivariate distances between different face identities, and zeros indicated distance between same face identities). PS showed overall lower fit with this identity model from approximately 172 ms to 746 ms (see supplementary figure 2, p<.05, cluster-corrected), indicating extended impairment in neural processing of face identities (Johnston et al., 2016; Wiese et al., 2019a; Yan & Rossion, 2020). Notably, this reduced distances between face-identity representations cannot be solely attribute to signal-to-noise differences in the brain of the patient; PS RDMs showed on par correlations with other models of categorisation (e.g. scenes vs. non-faces, objects vs. non-faces; see supplementary figure 2) compared to typical participants.

We also tested whether these impaired face-identity representations in PS could generalise to neural processes of individuals with typical abilities, i.e. a critical prediction in the quantitative (vs. qualitative) accounts of individual differences in face-recognition ability (Barton & Corrow, 2016; Bobak et al., 2017; Hendel et al., 2019; Maguire et al., 2003; Price & Friston, 2002; Rosenthal et al., 2017; Vogel et al., 2005; Zadelaar et al., 2019). We correlated the individual neural individuation score across time with face recognition ability within the control participants only. This showed that neural distance between face identities correlated positively with face specific performance scores across neurotypicals (peak-*r* = 0.5305, peak-*r*_time_ *=* 418 ms; *ps*<.05 from 400-434 ms). Similar results were obtained with face vs. non-face accuracy in the one-back task, or the CFMT+ taken separately.

Previous studies have shown that prosopagnosic patient PS displays somewhat typical face-selective activation (i.e. typically higher face vs. non-face BOLD activity) throughout most of the cortical face network (Gao et al., 2019). We asked whether fine-grained temporal multivariate analyses could reveal more subtle differences in the neural representations of PS. On top of face-identity, our various experimental conditions offered a mean to assess the putative typicality of PS’s neural response for important categorical distinctions (e.g. face-gender, face vs. non-face, scenes vs. non faces, etc.). We derived RDM correlations with these categorical models in an identical way than with the face-identity model. For example, face-gender information was assessed as the fit between brain RDMs and a face-gender binary model (where ones indicated multivariate neural distance between faces of different sex; see (Dobs et al., 2019). As can be assessed in supplementary figure 2, while PS showed impaired neural fit across face models (particularly with face-identity, left column), she showed overall typical fits with the non-face models (right-column), thus replicating the decoding analyses shown in **figure 2**.

**Supplementary figure 2.**
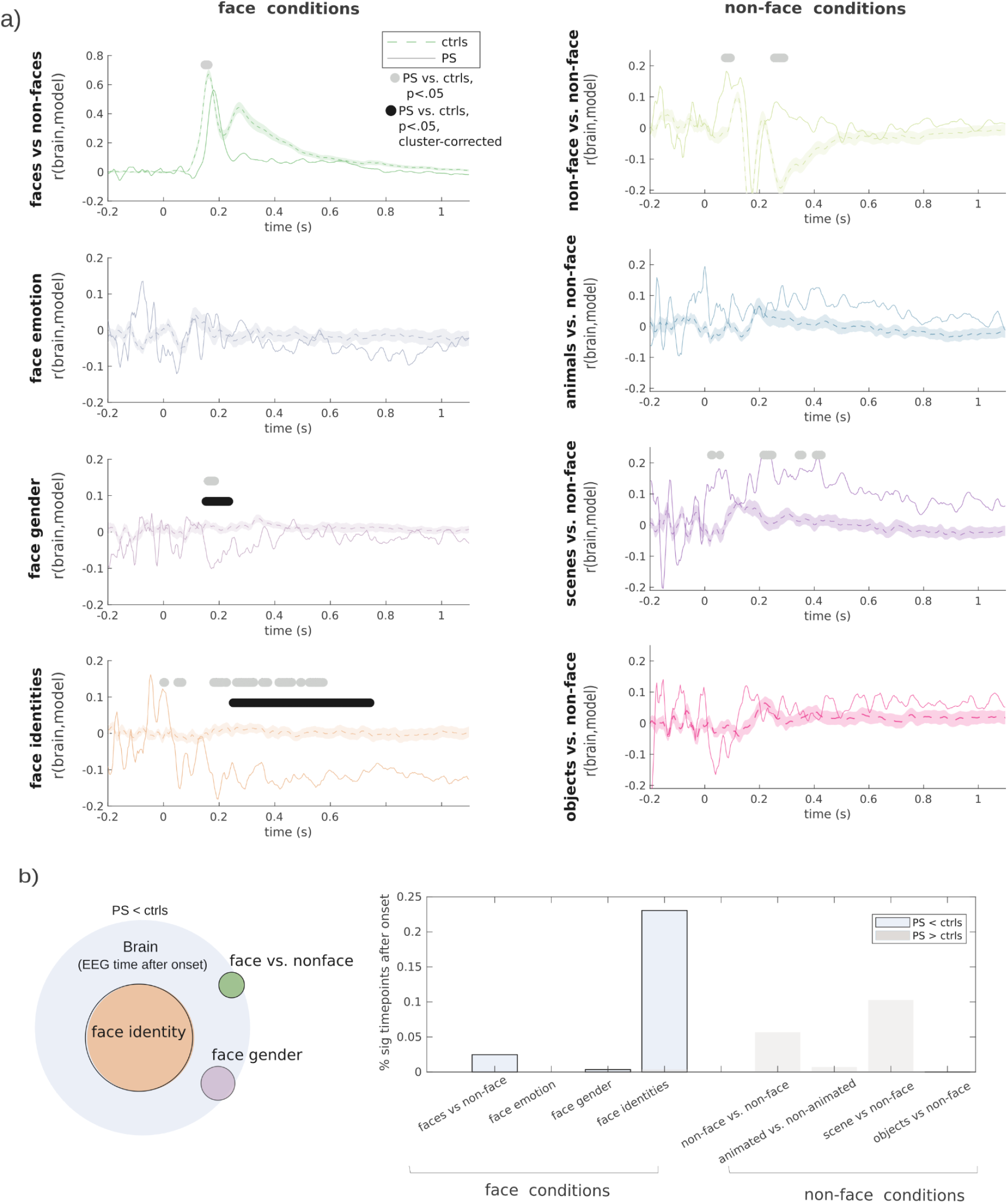
Time course of categorical distinctions compared between PS and neurotypicals. Brain RDMs for PS and controls were correlated at every 4 ms step with categorical models of face conditions (e.g., female vs. male faces in face gender condition) and non-face conditions (e.g., scenes vs. non-faces). Comparison of these correlations between PS and controls showed significant contrasts (Howell-Crawford t-tests, *p*<.05, coloured points). These effects are summarized in panel b for significant contrasts where PS showed reduced fit compared to controls. PS showed reduced neural fit exclusively with face model conditions, while she showed better fit with scenes vs. non-face model conditions. Proportion of all time points after EEG onset showing significant contrasts (PS < controls, *p*<.05, uncorrected) is shown on a Venn diagram as a quantitative summary of PS’s impaired brain computations for face identity (orange), face vs. non-face (green) and face-gender evidence (purple). The total surface area for the “brain” section represents all possible time points after onset (0-1100 ms in 4 ms steps), while the surface of each category represents a portion of these time points. Overlap indicates significance over the same time points.

**Supplementary figure 3.**
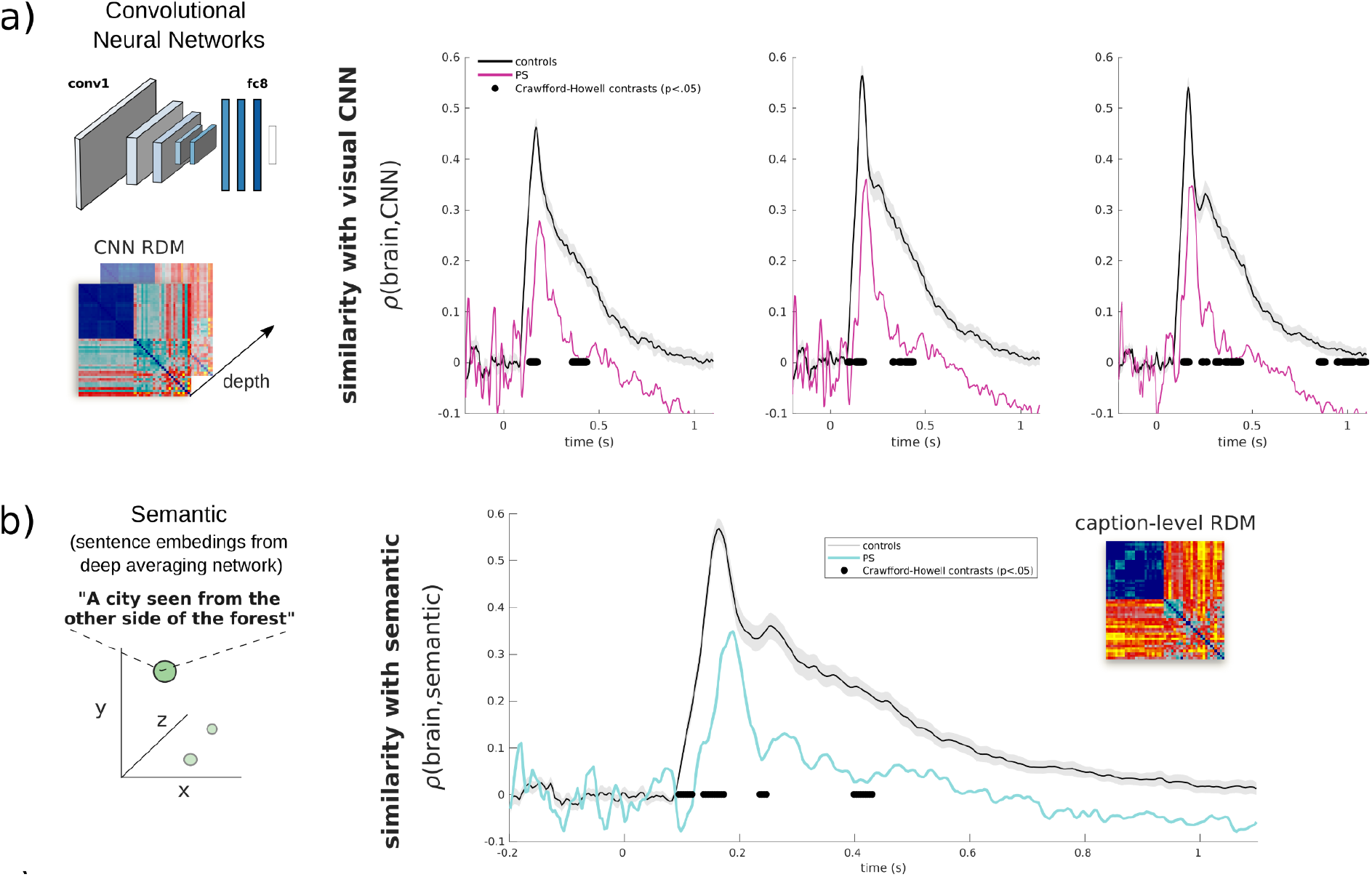
Time course of visual and semantic computations, unconstrained. **a**) Time course of correlation between brain RDMs and AlexNet RDMs is shown for PS (coloured curve) and controls (grey curve). Each column shows correlations with different layer RDMs 2, 4, and 6 in ascending order. We found overall lower similarity of visual computations within the brain of PS compared to controls (black dots indicates significant contrasts, Howell-Crawford t-tests, *p*<.05), indicating similar results to those observed when removing the shared information between CNN and the semantic (caption-level) model, in **figure 4**). Time course of correlation with RDMs of the semantic model was also significantly lower in the brain of PS compared to controls (cyan curve; black dots indicate significant contrasts, *p*<.05). The shaded areas of all curves represent the standard error for the controls.

**Supplementary figure 4.**
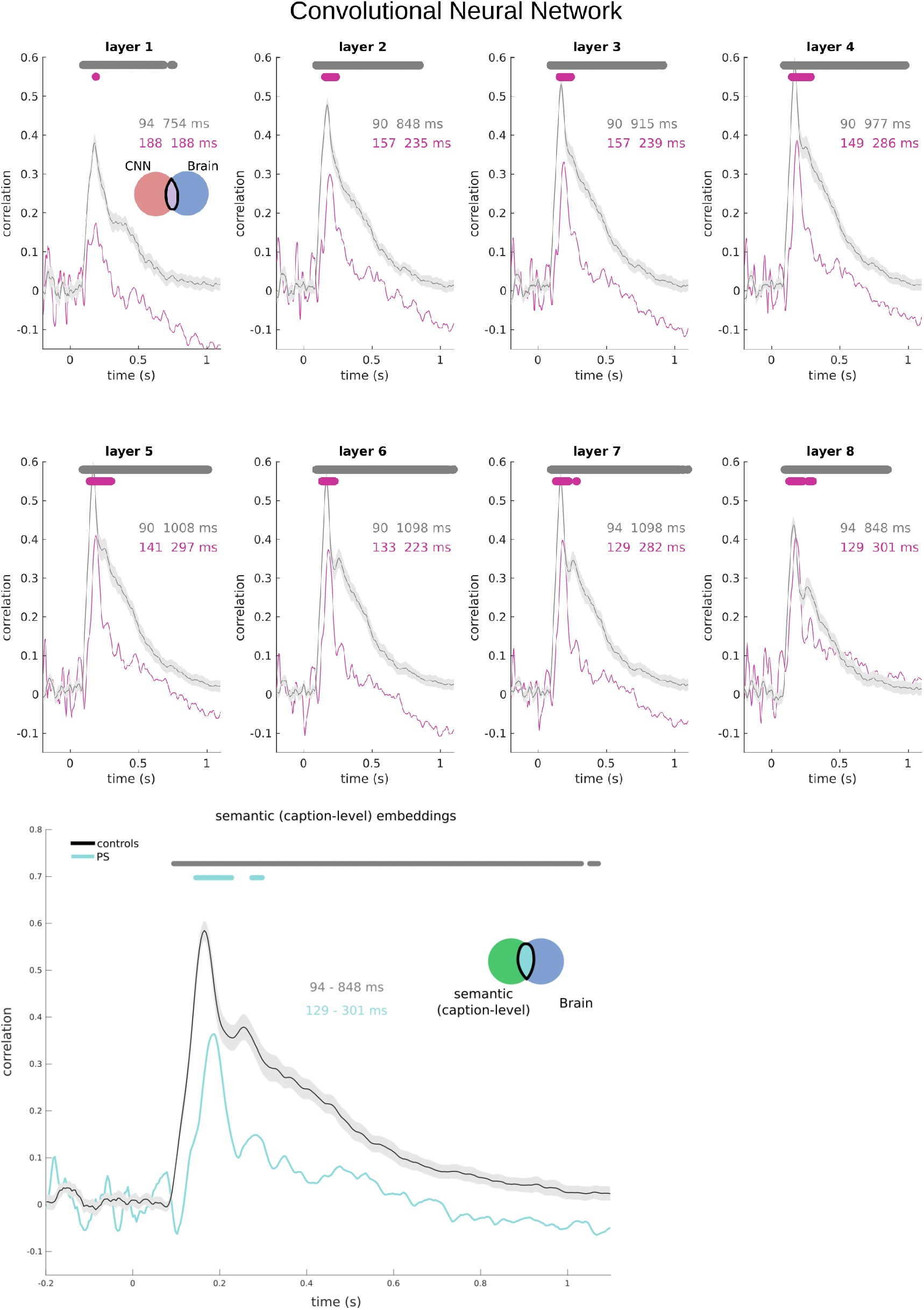
**a**) Time course of correlation between brain RDMs and AlexNet RDMs is shown for PS (coloured curve) and controls (grey curve) across all CNN layers. Each graph shows correlations with different layers, with significant time intervals indicated in the graph. **b)** Time course of correlation with RDMs of the semantic model is also shown for PS and controls. The shaded areas of all curves represent the standard error for the controls.

**Supplementary figure 5.**
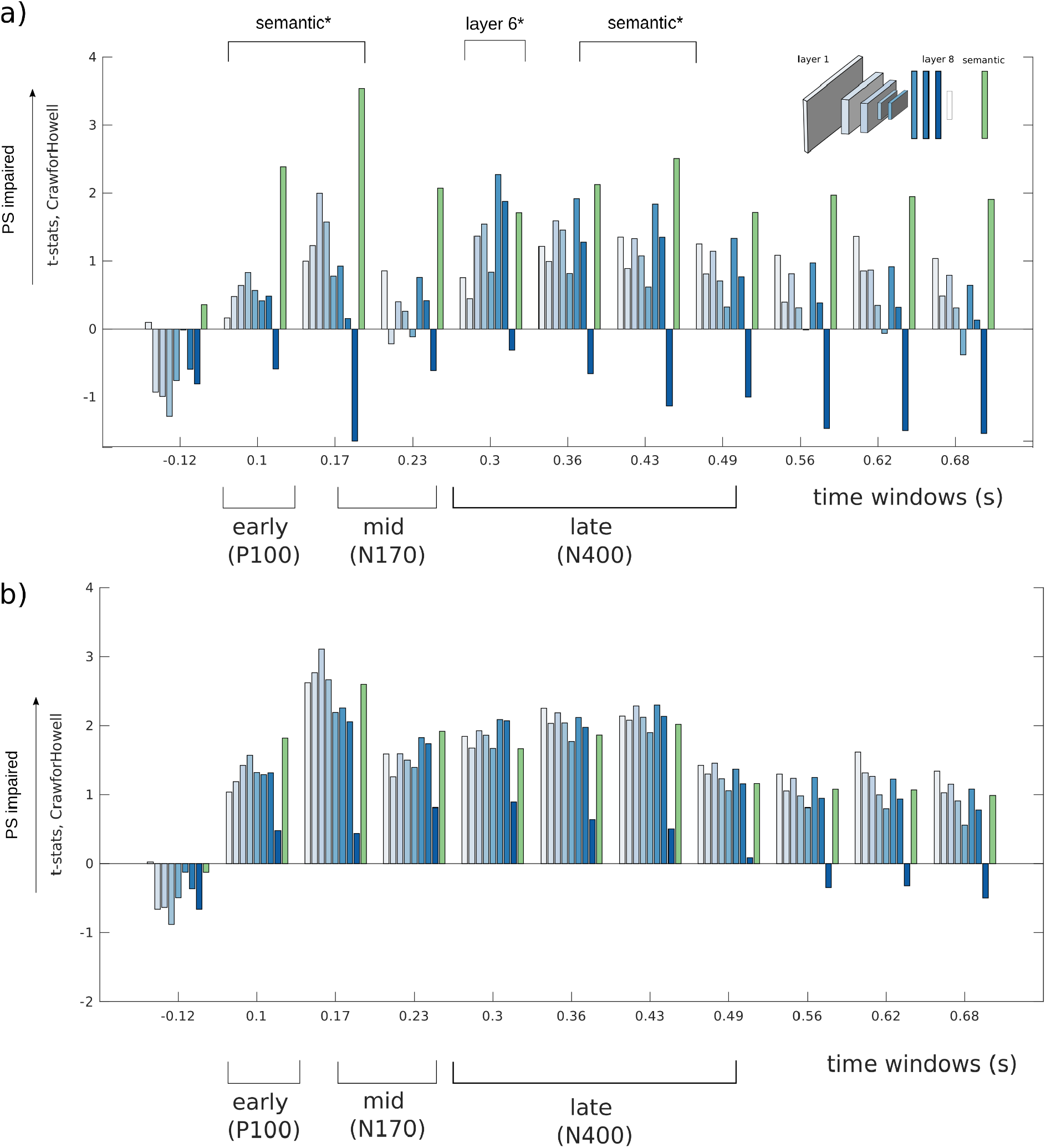
Time course of categorical distinctions compared between PS and neurotypicals. Correlations of brain RDMs to visual model (blue plots, with deeper blue corresponding to deeper CNN layers) and semantic model (green plot) was assessed in PS and controls for brain RDMs computed at every 60 ms window. **(a) (b)** Crawford-Howell t-statistics comparing controls vs. PS on these correlations are shown for the visual CNN (blue bars, with deeper blue corresponding to deeper CNN layers) and the semantic model (green bars; significant comparisons are indicated with an asterisk). T-tests comparing partial correlations between PS and controls are shown in **(a)** and those comparing simple correlations with the models are shown in **(b)**.

